# FreiBox: A versatile open-source behavioral setup for investigating the neuronal correlates of behavioral flexibility via 1-photon imaging in freely moving mice

**DOI:** 10.1101/2022.11.18.517059

**Authors:** Brice De La Crompe, Megan Schneck, Florian Steenbergen, Artur Schneider, Ilka Diester

## Abstract

To survive in a complex and changing environment, animals must adapt their behavior. This ability is called behavioral flexibility and is classically evaluated by a reversal learning paradigm. During such a paradigm, the animals adapt their behavior according to a change of the reward contingencies. To study these complex cognitive functions (from outcome evaluation to motor adaptation), we developed a versatile, low-cost, open-source platform, allowing us to investigate the neuronal correlates of behavioral flexibility with 1-photon calcium imaging. This platform consists of FreiBox, a novel low-cost Arduino behavioral setup, as well as further open-source tools which we developed and integrated into our framework. FreiBox is controlled by a custom Python interface and integrates a new licking sensor (Strain Gauge lickometer) for controlling spatial licking behavioral tasks. In addition to allowing both discriminative and serial reversal learning, the Arduino can track mouse licking behavior in real time to control task events in a sub-millisecond timescale. To complete our setup, we also developed and validated an affordable commutator, crucial for recording calcium imaging with the Miniscope V4 in freely moving mice. Further, we demonstrated that FreiBox can be associated with 1-photon imaging and other open-source initiatives (e.g., Open Ephys), to form a versatile platform for exploring the neuronal substrates of licking based behavioral flexibility in mice. The combination of the FreiBox behavioral setup and our low-cost commutator represents a highly competitive and complementary addition to the recently emerging battery of open-source initiatives.

**Significance Statement:** Behavioral flexibility is essential to survive in a complex and changing environment. To study this cognitive ability in freely-moving mice, we developed a versatile, low-cost, open-source behavioral setup, called FreiBox, allowing us to investigate the neuronal correlates of licking-based behavioral flexibility. FreiBox is controlled by a custom Python interface and integrates a new licking sensor for controlling spatial licking behavioral tasks (e.g. discriminative learning, reversal learning). We also developed and validated an active commutator to record calcium imaging with the Miniscope V4 in freely moving mice. Finally, we demonstrated that FreiBox can be associated with 1-photon imaging and other open-source initiatives, to form a versatile platform for exploring the neuronal substrates of licking based behavioral flexibility in mice.

## Introduction

Behavioral flexibility is essential to live in a complex and changing environment and to adapt to the infinite contexts of our daily life. An action which was appropriate in the past can become obsolete after a contextual change. This cognitive ability requires multiple cognitive processes which are supported by a vast cortico-subcortical network (Floresco & Jentsch, 2011; Izquierdo et al., 2017; Schoenbaum et al., 2009). By using behavioral flexibility paradigms, it is possible to explore cortical, basal ganglia, or thalamic networks activities, as well as to cover many fields in behavioral neuroscience (e.g., sensory-discrimination, decision-making). To do so in an animal model, we need to measure behavioral responses that can be attributed to association, contingency learning, decision, or adaptation.

Typical behavioral responses in freely moving animals are nose pokes (NP) in specific chambers, visits in maze arms, or presses on a specific lever to obtain a reward (Castañé et al., 2010; Izquierdo et al., 2017; Parker et al., 2016). However, no currently available setup permits measurement of directional licking, a behavioral paradigm which can access numerous cognitive processes in head-restrained mice, in a freely moving animal (Bollu et al., 2021; Catanese & Jaeger, 2021; Mayrhofer et al., 2019; Y. Wang et al., 2021; Wu et al., 2020a). Focusing on licking behavior, we have designed a fully open-source and cost-efficient (~900 €) behavioral platform to explore behavioral flexibility in freely moving mice. This new box, called FreiBox (Fig. 1), is capable of monitoring a wide variety of licking-based tasks, such as discriminative learning (DL) and serial reversal learning (RL), by tracking online licks with strain-gauge-based force lickometers.

**Figure 1.**
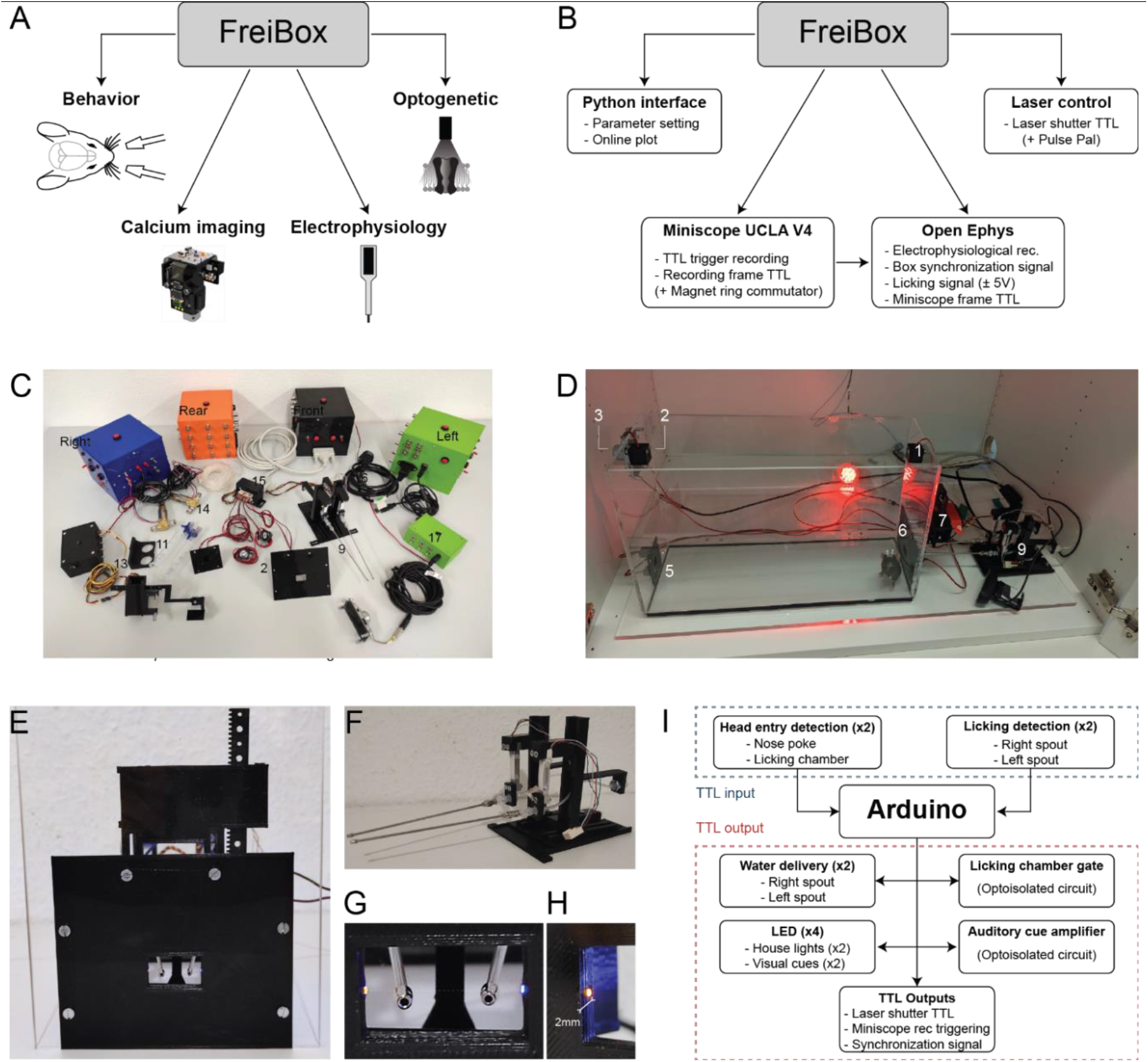
FreiBox platform and its operant instruments. **(A)** The FreiBox platform combines licking based behavior with one-photon calcium imaging, electrophysiological recordings, and optogenetic manipulation. **(B)** FreiBox is connected to a workstation equipped with a Python interface to set the task parameters and perform online analyses of the behavioral results. To perform optogenetic experiments, FreiBox can control a laser directly with an Arduino® or indirectly via a PulsePal device. An OpenEphys acquisition system integrated into the platform performs electrophysiological recordings and can be used to synchronize behavioral events and the frame signal of the Miniscope UCLA V4 register. An open-source and cost-efficient commutator has also been developed to perform calcium imaging (see Appendix 1). **(C-D)**. Pictures of the different components (C) assembled to build FreiBox (D). Component numbers: 1. and 2. house light; 3. speaker; 4. auditory tone generator; 5. nose poke (NP) chamber; 6. licking chamber (LC); 7. LC Gate; 8. LC gate controller; 9. strain gauge (SG) lickometer; 10. relay cable box; 11. water delivery pumps; 12. water tanks; 13. water tank holder; 14. water pipes; 15. D-SUB 25 cable; 16. power cable; 17. audio cable. **(E-H)**. Picture of the licking chamber (E) equipped with a gate, two lickometers (F) and a head entry detection module, based on the breaking of an IR beam (G) relayed by two optic fibers placed at 2mm from the entrance (H). **(I)**Input/Output transistor-transistor logic signal (TTL) mapping received and sent by the Arduino to react and control the electronic modules. In response to a head entry in the NP/LC or licking on the spouts, the Arduino sends a TTL pulse to deliver water, turn on and off the house lights, open the LC gate, generate an auditory cue, generate a synchronization signal, and control other devices (e.g., a laser shutter).

To track licking behavior online, conventional approaches use capacitive or optical methods to isolate individual licks. However, these methods generate electrical noise or are difficult to implement in freely moving mice. To address these issues, we developed a unique noiseless and sensitive mouse force lickometer ideally suited for freely moving conditions. To facilitate the integration of this new device into our platform, we added an adjustable online threshold detection system, making it compatible with nearly any behavioral platform, including those for head-fixed preparations. In addition, we designed a new licking chamber equipped with a gate, to control the accessibility of the licking spout, and a fiber-optic-based infrared beam system with a very short sensing distance (2 mm) to facilitate its application to detect NP with implanted animals. We showed that by combining FreiBox with other open-source technologies, we are able to perform one-photon calcium imaging in freely moving mice for a competitive cost per unit (<9 k€). To this end, we have also included a low-cost (<150 €) commutator that is equipped with a novel sensing system that improves online tracking of cable rotations without impairing the one-photon calcium imaging via a miniscope. Finally, to support the open-source philosophy, we built FreiBox by assembling multiple modules that are shared and fully available on our lab’s Github (https://github.com/Optophys-Lab/FreiBox).

## Methods

### FreiBox platform and its operant instruments

Influenced by the lever-press tasks in which an animal chooses between pressing on the right or left lever (Boulougouris et al., 2007; Brady & Floresco, 2015), FreiBox (Fig. 1C,D) offers the mice directional licking. We equipped FreiBox with a NP and a licking chamber (LC) placed at opposing sides of the Plexiglas box (40 × 20 × 15 cm) (Fig. 1C-H). The LC contains two strain gauge (SG) lickometers and a chamber gate (Fig. 1E-G). The LC gate provides control over the licking access, similar to the lever retraction in commercial boxes. Inspired by previous optical lickometers (Isett et al., 2018; Schoenbaum et al., 2001), we included an optic fiber infrared (IR) detection system at 2 mm from the entrance of both chambers to detect head entries (Fig. 1G,H). This design offers the advantage of decreasing the detection distance from the entrance and reducing the risk for the animal to touch the box walls with its implants. In addition to the operant instruments, we added several components (Fig. 1C,D) that can be used to deliver water or to generate light or auditory cues. An Ultimaker 3 printer equipped with black PLA filament was used to make the 3D-printed parts.

To control FreiBox and its electronic modules, we used the programmable microcontroller Arduino Mega 2560 (Arduino^®^, Fig. 1I). This board can be easily interfaced with a Python library (Pyserial) to control behavioral tasks, set the task parameters, or collect results to perform online plotting. Although this Arduino board has the major advantage of interacting directly with sensors or other electronic modules (e.g., sound card) through several analog and digital Input/Output (I/O) pins and multiple libraries (Fig. 1I), it is a “one core” device which can execute only one instruction at a time. This dramatically reduces the processing speed. According to the manufacturer (https://www.arduino.cc/reference/en/language/functions/analog-io/analogread/), the analog sampling rate is relatively slow (<10 kHz for a single channel) and decreases with the number of recorded channels. This contrasts strongly with the high-speed capability of an Arduino to read and write digital signals with the Arduino library ‘DIO2’ (≈ 4 μs, 250 kHz, https://www.arduino.cc/reference/en/libraries/dio2/). The Input/Output reactivity of the Arduino Mega 2560 (Extended Data Fig. 1-1) has a delay of less than 20 μs. Based on those results, we built electrical modules which interact with the Arduino board by using digital signals only (Fig. 1I). This transistor-transistor logic (TTL)-based modular organization facilitates the electronic integration and transposition into different behavioral controller platforms (Raspberry pi or Teensy). Hence, we developed several hardware modules which can be incorporated or removed depending on the task needs. We provide here the circuits used to build FreiBox (Extended Data Fig. 1–2).

### Sensitivity evaluation of the strain gauge lickometer

When a mouse licks on the spout of the SG-lickometer (Fig. 2A,B), the amplified SG-signal (SGs) (Extended Data Fig. 1–2A-C) exhibits large peaks (Fig. 2C and Extended Data Fig. 2–1A) which can be easily recorded by any analog-to-digital converter (e.g., OpenEphys), and analyzed offline (see the following section) to extract individual licks (Fig. 2C, ‘SG-offline detection’). Given the large signal-to-noise ratio of the SGs, we connected a voltage comparator circuit (Extended Data Fig. 1–2C) to track the licking behavior online (Fig. 2C, ‘SG-online TTL’). Such a circuit generates a TTL pulse when the lick peaks of the SGs cross a defined reference voltage threshold. For the validation experiment described below, this threshold was set at 100 mV above the baseline and kept constant for all mice.

**Figure 2.**
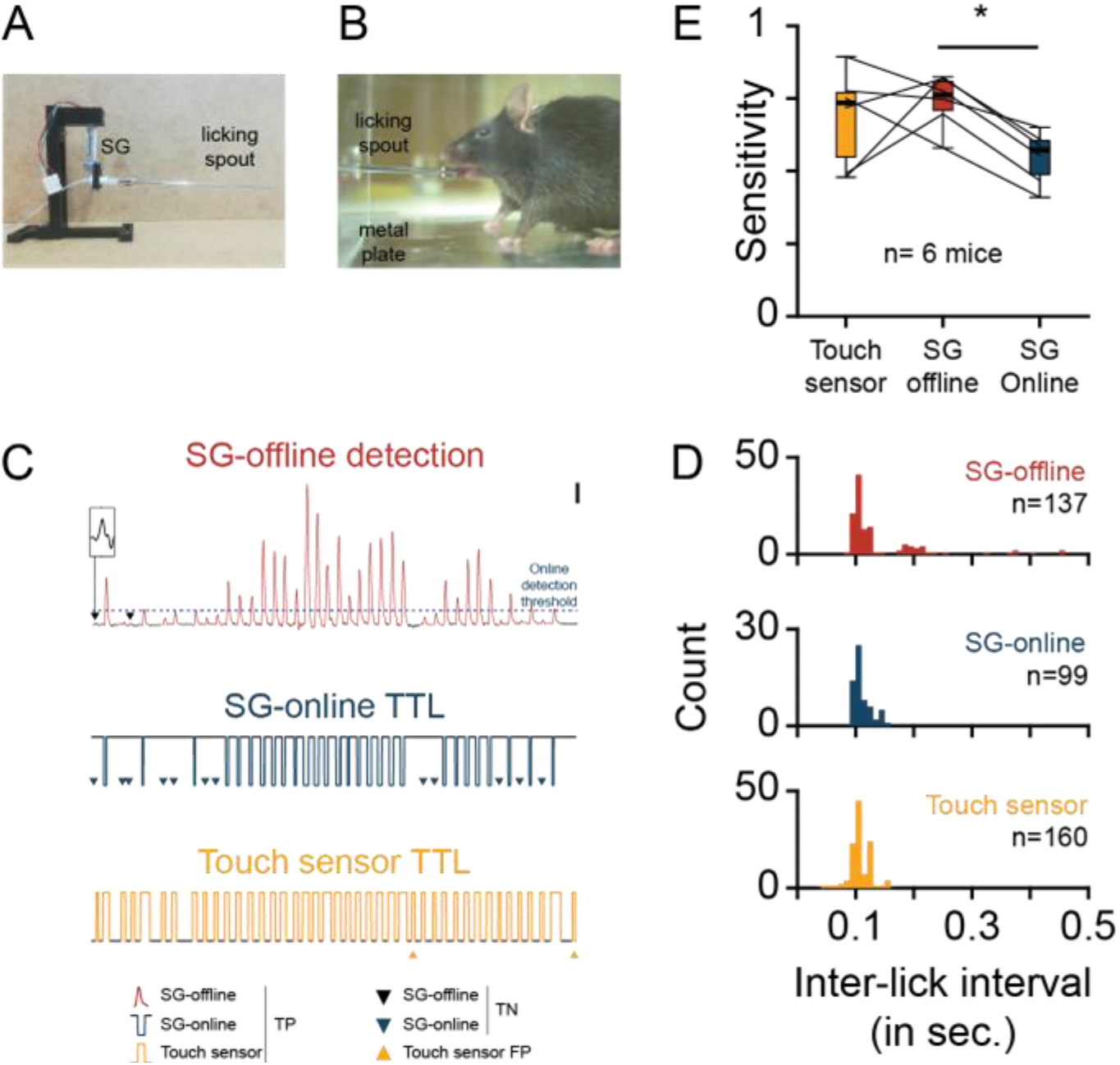
Validation of the strain-gauge lickometer. **(A)**. Picture of the SG lickometer, composed by a 3D-printed holder attaching a SG connected to a licking spout. **(B)**. Picture taken by the high-speed camera used to define the ground truth licks and compare the sensitivity of the SG and touching lickometers. The metal plate on the box floor was used to ground the mice when the animal tongue touches the metallic spout. **(C)**. 5 s example of true positive (TP), false negative (FN) and false positive (FP) licks from the 3 licking detection methods (SG-offline, SG-online and touch sensor), defined by comparison with the ground truth licks defined by a high-speed video recording. On the top trace (scales: 5 s, 100 mV; Inset scales: 200 ms, 50 mV), the online detection threshold used to induce the SG-online TTL is plotted. **(D)**. Distribution histogram of the inter-lick interval of the true positive licks recorded with the 3 licking methods shown in C. **(E)**. Box-and-whisker plots comparing the sensitivity of touching sensor and SG lickometers. one-way Anova repeated measures (F=4.906, *p*=0.033) followed by a Tukey test (q=34.3), \**p*=0.031.

To test whether the sensitivity of the SG-lickometer (Fig. 2) is comparable to the classical touch sensor (TS), we placed 6 water-restricted mice in a water-delivery chamber (20 cm × 20 cm × 20 cm), containing a single licking spout attached to a SG (Fig. 2A) and connected to the TS circuit (Goltstein et al., 2018). The box floor was covered with a metal plate connected to the ground of the TS circuit. High-speed video recordings (1000 frame per second, Basler ace camera, acA1300-200uc) were simultaneously performed (Fig. 2B) to control the contact of the mouse’s tongue with the spout and measure the ‘ground truth licks’ (GT-licks). The videos were then annotated offline with the software BORIS (Behavioral Observation Research Interactive Software) to extract the licks’ timestamps. In order to align the licking detection timestamps, the TTL from the camera (indicating the exposure window of each video frame) were recorded with an Open Ephys system (sampling rate: 30 kHz), simultaneously with the SGs, SG-online TTL, and TTL of the TS. The GT-licks were then aligned to the SGs (analog and SG-online TTL) and the TS TTL, to count true positive (TP) and false positive (FP) licks (Fig. 2C). The sensitivities (Sensitivity = [TP / (TP + FN)]) and positive predictive values (PPV = [TP / (TP + FP)]) were evaluated to compare the detectors.

To extract individual licks, the SGs has been first down-sampled (with an antialiasing low-pass filter) to 1 kHz (Matlab^®^ function “resample”) and low-pass filtered at 64 Hz. The function “findpeak” (Matlab®) was then used to extract the peaks of individual licks. To improve the detection, we used the following parameters: a minimal peak distance of 60 ms, a minimal prominence of 0.015, and a minimal peak width of 50 ms. We applied a matching procedure to validate each detected lick, by comparing the delay of each detected lick with its closer GT value. A lick was considered as TP if this delay was less than the minimal inter-lick interval used for the SG-offline detection (60 ms).

### Animals and water restriction protocol

All animal procedures were performed in accordance with the guideline RL 2010 63 EU and approved by the Regierungspräsidium Freiburg. The animals were housed (Blueline Type 1284 L, Tecniplast) with a humidity between 45 and 65% and a temperature between 20 and 24 degrees, under a 12 h light-dark cycle (light period from 8 AM to 8 PM). Mice received food ad libitum. A total of 11 mice (7-10 weeks, male) were used in this study: 8 C57BL/6J were used to validate the intra-session RL; 1 C57BL/6J and 2 Thy1-GCaMP6f (C57BL/6J-Tg(Thy1-GCaMP6f)GPS.5.17 DKim/J) were used to evaluate the frame loss percentage of the commutator and to explore the calcium activity of the orbitofrontal cortex (OFC) during DC.

To motivate the mice to perform the behavioral task, they were maintained under water restriction up to 5 days of the week followed by 1-2 days of water ad libitum. During the water restriction period, a careful monitoring of the mice weights was performed on daily basis. Care was taken to ensure that they did not weigh less than 80% of their free-feeding weight measured at the start of the week of deprivation. For training days, the animals consumed at least 1 ml of water. When weight fell below 80%, additional water was given until they recovered (>80% of the initial weight). In the periods without behavioral assays, the mice received water ad libitum.

### Behavioral task (244 words)

The RL task is a freely moving directional licking paradigm (Fig. 3A-C) adapted from a head-restraint condition (Mayrhofer et al., 2019). During the DL phase, the mice had to explore to find the reward spout by licking on one of the two available spouts (Fig. 3A). After reaching an online performance criterion (70% hit in a sliding average window of 15 trials), the reward spout is automatically changed and the mice need to adapt their behavior (Fig. 3A, RL phase). To initiate a trial and open the LC gate, the mice must perform a NP (Fig. 3B). They can then walk (walking delay, WD) to enter their head into the LC (head entry, HE) and lick on the correct spout during the licking response period (LR, 2 sec). At the end of the LR, an auditory cue is played, consisting of a pure 5 kHz tone for the correct licking trials (or hit trials) or white noise for missed trials (entry in the LC without licking on a spout) or error trials (licking on the incorrect one). After an incorrect or missed trial, the gate is automatically closed. In contrast, a reward anticipation period (RA) is maintained 1 s before the water delivery for the hit trials, to allow recording of reward-related activity prediction. In this configuration the auditory cues become predictive of the outcomes. Finally, the LC gate is closed and the trial finished after 5 s post-reward (PR) interval, allowing reward consumption.

**Figure 3.**
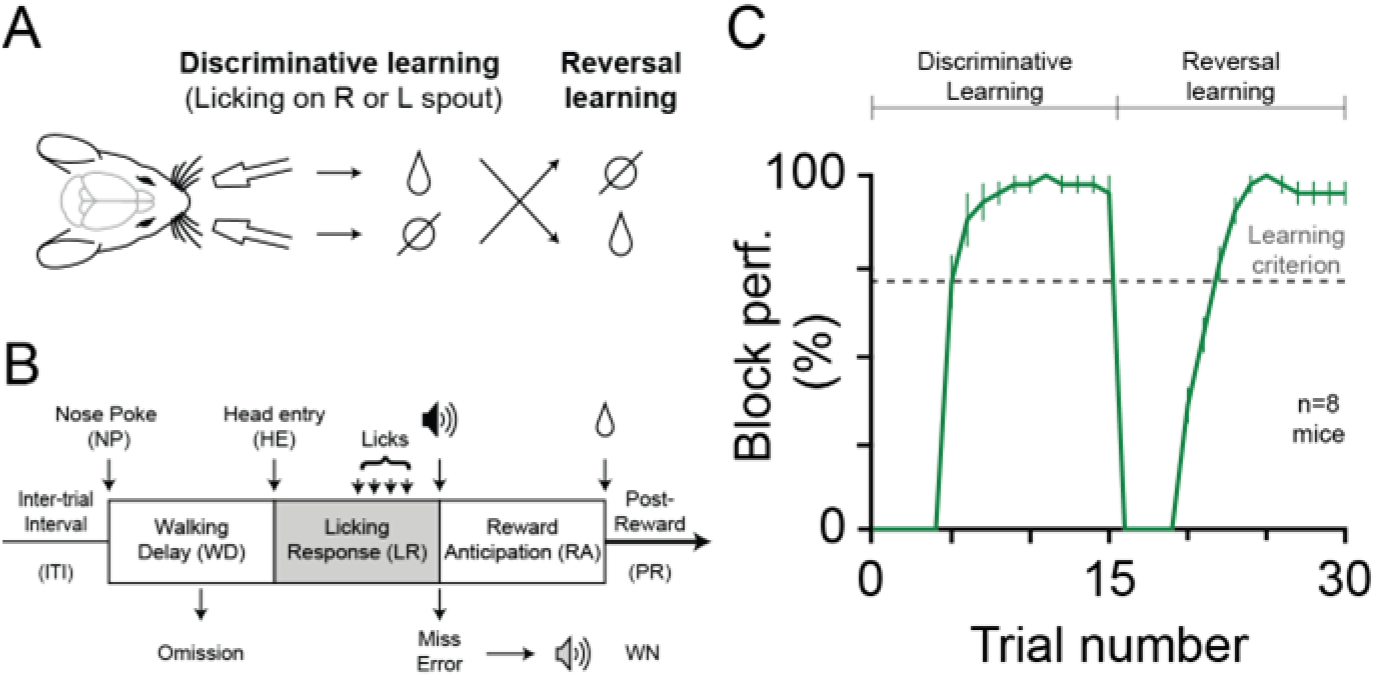
FreiBox control of intra-session reversal learning. **(A-B)**. Timeline (A) and description of the dual licking spouts intra-session RL task (B). **(C)**. Population averaging (n=8 mice) of the block performance during RL (mean ± SEM). For each expert animal, the best session was selected to perform averaging. The block performance was calculated with a moving average of 5 trials.

### Behavioral training

To train each mouse to perform the task, we slowly introduced each task component and personalized the training to respect the individual learning rate. The first stage consisted of the association between the licking spouts, the water reward, and the auditory cues. To accomplish this, the LC gate was left opened until the mouse placed its head in the LC. If the mouse licked on the spout, the pure tone cue was played immediately, the reward delivered, and the gate closed after 4 s. In contrast, if the mouse did not lick on the spout 2 s after the head entry (missed trial), the white noise cue was played and the gate closed. At this stage, until stage three, only one spout was presented in the LC and a session for each licking spout was programmed to avoid a spatial bias. The second stage was similar, except that the licking response and anticipatory response were introduced. In the third stage, the NP was introduced, meaning that the LC gate was kept closed until the mouse performed a NP. During the fourth stage, we introduced the DL by presenting both spouts in the LC. To help the animal, we used immediate feedback by playing the white noise cue immediately after the mouse licked on the wrong spout, followed by LC gate closing. In the next training stage, the final paradigm was used to perform inter-session reversal of DL session. In such configuration, the reward spout learned during DL in the morning is reversed for the afternoon sessions. Finally, the intra-session reversal task was used. At this final stage, only one session per day was used to maintain a high motivation level in the animals. With this protocol, the mice learned the task in a few weeks.

### Surgery

All animal procedures were performed in accordance with the guideline RL 2010 63 EU and approved by the Regierungspräsidium Freiburg. General anesthesia was performed using a mixture of oxygen and isoflurane (3-5% induction, 1.5-3% maintenance, CP Pharma), associated with a subcutaneous buprenorphine injection (0.05 mg/kg, Temgesic^®^). The mice were then fixed on a stereotaxic frame (Model 942, Kopf^®^) and placed on heating blanket (Rodent Warmer X1, Stoelting^®^). During surgery, the depth of anesthesia was determined by the extinction of the pain reflexes (inter-toe and eyelid closure reflexes), and the eyes were protected from dehydration with eye ointment (Bepanthen, 798-037, Henry Schein). Five minutes before the rostro-caudal skin incision, a local surface lidocaine anaesthesia (xylocaine gel 2%, 1138060, Shop-Apotheke) was applied to the scalp. To maintain the fluid balance, supplementation with 1 mL of Ringer solution was performed every 2 h.

For 1-photon calcium imaging in C57BL/6J mouse, a virus injection procedure (De La Crompe et al., 2020) was performed right before gradient-index (GRIN) lens implantation. After craniotomy, a virus solution (rAAV1-hSyn-jGCamP7f, 104488-AAV1, Addgene) was injected under stereotaxic conditions (400 nL, 3.10^12^ viral genomes/mL) into OFC (AP: +2.5 mm from Bregma, ML: +1.5 mm from midline, DV: −1.8 mm from cortical surface) using glass microcapillary (tip diameter ≃35 μm; 1–5 μL Hirschmann® microcapillary pipette, Z611239, Sigma-Aldrich) connected to a custom Openspritzer pressure system (Forman et al., 2017). To perform GRIN lens implantation, we followed the instructions provided by the Miniscope V4 group (http://miniscope.org/index.php/Online_Workshop). After craniotomy and a tunnelling procedure using 25 G needle, the GRIN lens (diameter: 0.5 mm; pitch: 2; 1050-002183, Inscopix®), pre-assembled to the baseplate, was inserted above the OFC (AP: +2.3 mm from Bregma, ML: +1.5 mm from midline, DV: −1.8 mm from cortical surface). The lens was then secured by using a light-blocking cement obtained by mixing a small amount of a black pigment (black iron oxide, 832036, Kunstpark) with a dental cement (Paladur, Kulzer GmbH, Hanau, Germany). To build the mounted GRIN lens and adjust the working distance, we used a custom miniscope holder (https://github.com/Optophys-Lab/FreiBox) in combination with a lens holder available on the Moorman’s Lab github (https://github.com/moormanlab/miniscope-goodies) before cementing the GRIN lens to the baseplate. At the end of surgery, a cap was secured to the baseplate to protect the lens. All miniscope materials have been purchased at OpenEphys store or 3D-printed with a Ultimaker 3 printer and black PLA filament.

To prevent post-operative pain, we performed subcutaneous injections of buprenorphine (0.05 mg/kg every 6 h, Temgesic^®^) and carprofen (5 mg/Kg every 24 h, Rimadyl^®^, Zoetis) during the next 3 days post-surgery. To continue the analgesic treatment during the night, we provided a buprenorphine solution (0.1 g/L, Temgesic^®^) mixed in drinking water containing 5% (w/v) D-Glucose (G8769-100ml, Sigma Aldrich) to mask the drug taste. The weight and the general condition of the animals were monitored daily in the four days following surgery and weekly after the recovery of the animal.

### Miniscope recording and data analysis

Before starting the miniscope recordings, trained animals were habituated to hold a dummy 3D-printed miniscope during two to three sessions (https://github.com/Aharoni-Lab/Miniscope-V4). Once habituated, the mice were placed in the behavioral box with the miniscope secured to the implanted baseplate. During the behavioral sessions, Open Ephys recordings (sampling rate: 30 kHz) were performed simultaneously with calcium imaging to synchronize, with the same clock system, the data streams coming from the FreiBox (Fig. 4C, ‘Synchronization signal’) and the Miniscope V4 (Fig. 4C, ‘Miniscope frames’). After the acquisition (Fujitsu laptop, LIFEBOOK U749), the videos were preprocessed to remove the noise produced by the electrowetting lens driver (https://github.com/Aharoni-Lab/Miniscope-V4/wiki/Removing-Horizontal-Noise-from-Recordings) and cropped to preserve the part of the FOV containing only the GRIN lens. We then used the open-source tool library CaImAn (gSig/gSiz: 3/13; Giovannucci et al. 2019) to extract the spatial footprints and their associated temporal calcium traces, expressed as denoised temporal traces (parameter C of the source extraction algorithm CNMF-E). All the video processing steps were made with Python, and the following analyses were done in Matlab. To find behavioral correlations of the calcium traces, we first aligned all behavioral events to the closest recorded frames timestamp. We then calculated the waveform average of the median absolute deviation (MAD) of the denoised temporal traces. Based on this approach, we defined units as ‘modulated’ when their averaged calcium peaks crossed a 3xMAD threshold. Finally, we classified modulated footprints as initiation, licking, or reward neurons if their averaged MAD peaks occurred during the walking delay, the licking response, or reward anticipation period respectively.

**Figure 4.**
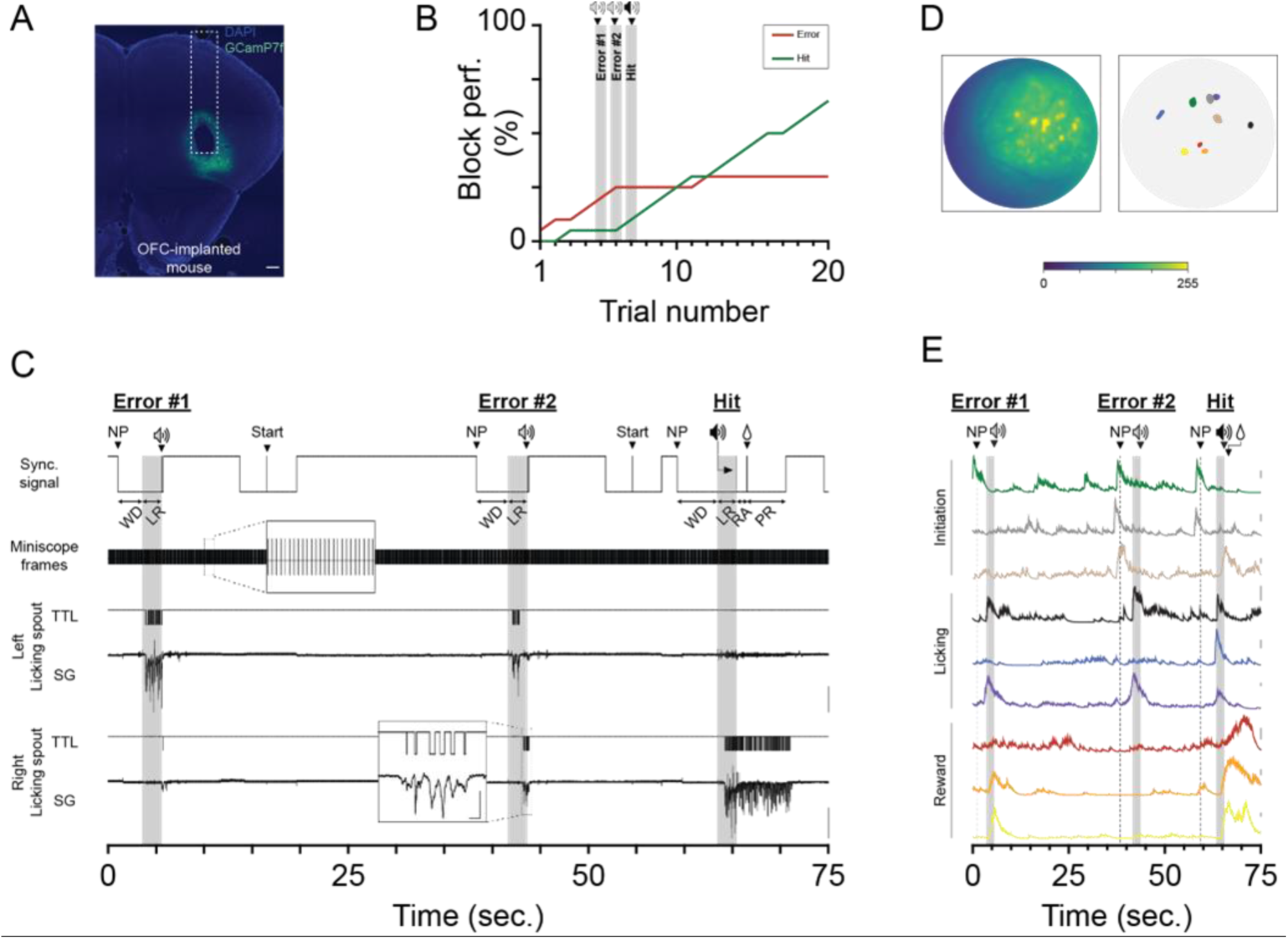
Combining one-photon imaging of OFC with FreiBox. **(A)**. GCaMP7f expression and GRIN lens track in OFC (scale: 250μm). **(B)**. Block performance of the implanted mouse (A) during the discriminative learning. The 3 highlighted trials are the same as shown in C and E. **(C)**. Simultaneous recording of the Miniscope and FreiBox signals used to synchronize calcium activity with behavioral events. **(D-E)**. Examples of 9 modulated neurons (D) and their calcium traces (E) from the video recorded in (C) during the same trial sequence as shown in (B). (D left) Maximum projection picture obtained on the first 10 k frames of the recording session. The spatial footprints (D right) and the calcium traces (E) were obtained with the open-source pipeline CalmAn (gSig/gSiz: 3/13). The calcium traces represent the denoised temporal traces (CalmAn parameter C, scales: 10 dF) and were colored in accordance with their spatial footprints in (D).

### Transcardiac perfusion and histological control

After completion of the experiments, the mice received an overdose of a pentobarbital sodium (1 mL, 1.6 g/L, i.p.; Narcoren). After respiratory arrest, a transcardiac perfusion (PBS 0.01 mM followed by PBS 0.01 mM/formaldehyde 4%) was performed for histological validations. The brain was kept overnight in a solution of PBS 0.01 mM (70011-051, life technologies)/formaldehyde 4% (v/v; E15711, Science Services), cryoprotected several days in a solution of PBS 0.01 mM/sucrose 30% (w/v, 1076511000, Merck Millipore), and then cut into 50 μm slices with a sliding microtome (SM2010 R, Leica Biosystems). The images were acquired with an Axioplan 2 microscope (Zeiss) controlled by the software Axiovision (v4.8, Zeiss). Shading correction and stitching were done with Fiji (based on ImageJ from NIH, USA) using the BaSiC (Peng et al., 2017) and Grid/Collection plugins, respectively.

### Figure design and statistical analysis

All plots were generated with Matlab and Python and were assembled with illustrator CS6 (Adobe) to generate the figures. For Fig. 3A and Extended Data Fig. 3–1D, the mouse head drawing was downloaded from https://scidraw.io/. The statistical analyses were performed (De La Crompe et al., 2020) using SigmaPlot 12 (Systat Software). For independent samples, we applied the normality (Shapiro-Wilk test) and equal variance tests. A t-test was used to compare samples if they were normally distributed and their group variances equal. Otherwise, the Mann-Whitney signed rank test was performed. For dependent samples, the paired t-test was used for normally distributed paired samples. In contrast, when the normality distribution test failed (Shapiro-Wilk test, *p*<0.05), the Wilcoxon signed rank test was performed. In same way, Friedman repeated measures ANOVA on ranks was used instead of one-way repeated measure ANOVA, when normality distribution was not verified. ANOVA was then followed by Tukey post hoc test.

## Results

### Validation of the strain gauge lickometer

Using licking as behavioral readout requires measurement of the force applied on the spout with a high sensitivity. Although it has been shown that SG-lickometers are able to detect licks when a mouse is drinking on a cup (Wang & Fowler, 1999), it has never been tested in a configuration where the SG is connected to a spout. Hence, before integrating our competitive SG-lickometer, we validated its sensitivity by comparing the SG-offline and SG-online methods with the well-established TS (Fig. 2D, ANOVA one-way repeated measure, n=6, F=4.906, *p*=0.033). In doing so, we found that the sensitivity of the SG-offline method is comparable to TS (TS vs SG-offline, 0.546 ± 0.0372 vs 0.740 ± 0.0374, Tukey test, TS vs SG-online, q=33.073, *p*=0.125) and significantly higher than SG-online (SG-online vs SG-offline, 0.685 ± 0.0676 vs 0.740 ± 0.0374, n=6, Tukey test, TS vs SG-online, q=34.30, *p*=0.031). In contrast, the TS and SG-online shared a similar sensitivity level (TS vs SG-offline, n=6, Tukey test, q=31.227, *p*=0.672). The PPV was not significatively different between the methods (Supplement S2D; TS vs SG-offline vs SG-online; ANOVA one-way repeated measure, n=6, F=0.166, *p*=0.850).

We tested whether the sensitivity difference between the offline and online SG methods can be compensated by lowering the value of the detection threshold. As illustrated in Fig. 2B and Extended Data Fig. 2–1A, the licks with a smaller amplitude did not cross the voltage threshold. By artificially adjusting the detection levels (Extended Data Fig. 2–1A,B), we showed that a reduction of only 20 mV is sufficient to improve the sensitivity greatly without compromising the PPV (Extended Data Fig. 2–1C,D). The potential to fine-tune the detection threshold greatly improves the real time detection performance. We implemented an Arduino oscilloscope (Extended Data Fig. 2–1E, modified from https://create.arduino.cc/projecthub/aimukhin/advanced-oscilloscope-955395), to monitor online the SG-signals and their thresholds. Taken together, these data demonstrate that the SG-lickometer can be successfully implemented to read out online licking behavior.

### Behavioral training and serial RL

After having validated the SG-lickometer’s ability to track individual licks, we programmed FreiBox to control an intra-session RL task (Fig. 3A,B). The results were obtained with a cohort of 8 mice trained during 4-6 weeks to acquire the intra-session reversal. As illustrated in their best (Fig. 3C) and the individual (Extended Data Fig. 3–1A,B) sessions, the mice were able to reach 70% of correct licking in a sliding average window of 5 trials, in fewer than 10 and 20 trials during the DL and RL blocks respectively. On average (Extended Data Fig. 3–1B), the mice required more trials to succeed in RL than in DL, a feature which has been described earlier (Castañé et al., 2010). Based on those positive results, we refined the FreiBox capability to control multiple task variants via an easy-to-use Arduino-Python GUI interface (Extended Data Fig. 3–1C). With such improvements, we were able to perform serial RL (Extended Data Fig. 3–1D) or probabilistic RL (Parker et al., 2016) by simply adjusting the block number or reward probability, respectively.

### Calcium imaging during discriminative learning

DL paradigms have often been used under 2-photon conditions to explore the role of cortical areas in decision making and motor execution (Chen et al., 2017; Galiñanes et al., 2018; Guo et al., 2017; Morrissette et al., 2019; Wu et al., 2020b). To test the feasibility of coupling calcium imaging in FreiBox with such a behavioral task, we developed an active miniscope commutator (Extended Data Fig. 4–1) and trained a mouse injected with the viral vector rAAV1-hSyn-jGCamP7f and implanted with a GRIN lens in OFC (Fig. 4A). Once the DL was complete, we performed recordings with the miniscope (Fig. 4B,C). To synchronize calcium data with behavior, we recorded simultaneously with an Open Ephys system, the analog and TTL SG licking signals, the miniscope frames TTL from the DAC box as well as the synchronization signal from the FreiBox (Fig. 4C). Fig. 4C shows an example of those signals recorded from two consecutive error trials (Error #1 and #2) followed by a hit attempt (Hit). That trial sequence was the turning point of the session because afterwards the hit performance continuously increased while the error rate stagnated (Fig. 4B).

To analyze calcium videos, we used the open-source tool library CaImAn (Giovannucci et al., 2019) to extract the spatial footprints (Fig. 4D) and their associated temporal calcium traces (Fig. 4E). After finding some calcium activities modulated during recorded behavior (see method), we aligned neuronal examples to behavioral events (Fig. 4E) and found that most of the modulated jGCamP7f-OFC expressing neurons can be separated into three main populations, depending on their peak activity in the task periods: initiation (during walking delay), licking (during licking response), or reward (during reward anticipation period). We confirmed this response distribution by repeating the same experiment in two trained Thy1-GCaMP6f mice during serial RL, as illustrated for one of them in Extended Data Fig 3–1D and Fig 4–1. These results illustrate the complex encoding of OFC neurons during goal-directed behavior and the ability of our task to dissect different neuronal processes.

## Discussion

Since the emergence of head-restraint preparations, directional licking has been used as a behavioral readout to access numerous cognitive processes including short-term memory (Y. Wang et al., 2021), decision-making (Wu et al., 2020b), or motor preparation (Chen et al., 2017). Conventional approaches for monitoring online licking behavior use electrical and optical sensors (Weijnen, 1998; Williams et al., 2018) or high-speed video recordings combined with deep-learning approaches (Bollu et al., 2021; Catanese & Jaeger, 2021). In general, while those lickometers are very efficient to detect the occurrence of individual licks, they are difficult to implement for online applications. For example, the gold standard TS lickometer requires proper grounding of the animal in order to detect licks while avoiding electrical noise (https://www.janelia.org/open-science/comparator-dual-lick-port-detector). To overcome those limitations, we developed and validated a cost-effective and sensitive force lick sensor which can be used with freely moving mice. In contrast to the available force lickometer which measures licking on a flat disk (Rudisch et al., 2022; Wang & Fowler, 1999), our SG-lickometer can be easily integrated to measure directional licking in a large number of behavioral setups, including head-fixed settings.

Based on our novel FreiBox platform, we are able to combine freely moving licking-based behaviors (e.g., DL, RL, or set-shifting) with the most recent optogenetic approaches. As a proof of concept, we performed miniscope calcium imaging in mice performing DL and RL. We recorded several response patterns in the OFC, demonstrating that our platform is able to capture complex neuronal dynamics reflecting several aspects of behavioral flexibility such as decision making, motor planning, and action evaluation. We recorded calcium activity related to nose-poking and licking, but also to a neuronal population expressing a long-lasting plateau following reward delivery. These findings corroborate results observed with electrophysiological striatal and cortical recordings of monkeys performing a RL task (Histed et al., 2009; Pasupathy & Miller, 2005). Altogether, these data introduce FreiBox as a useful tool to explore the neuronal substrates of behavioral flexibility underlying directional licking in rodents.

Adaptation of the methods to experimental questions is a major advantage of developing open-source tools (White et al., 2019). Our development of FreiBox and its low-cost parts represent complementary additions to the recently emerging battery of open-source techniques (White et al., 2019). Recently, open-source initiatives such as miniscope (Aharoni et al., 2019) and Open Ephys (Siegle et al., 2017) have opened the gate to the standardization of flexible, low-cost, and open-source technologies. In a global context of budget reduction, open-source projects and publications constitute a great advance to reduce the financial dependency of the laboratories without compromising research quality.

**Table 1.** Price list for assembling the FreiBox platform.

| ID | Name | Price (€) |
| --- | --- | --- |
| 1 | FreiBox | 903 |
| 2 | Commutator | 111 |
| 3 | Tools | 160 |
| 4 | Miniscope UCLA v4 | 1580 |
| 5 | Open Ephys system | 3479 |
| 6 | Laser Controler | 900 |
| 7 | Computer | 1500 |
| Total |  | 8633 |

## Acknowledgements

We thank David Eriksson for helpful remarks and advices and Christine Zeschnigk and Sabine Weber for technical support.

## Abbreviations

AP: Anteroposterior
DL: Discriminative learning
DV: Dorsoventral
FP: False positive
GRIN: Gradient-index
GUI: Graphical user interface
GT licks: Ground truth licks
IR: infrared
I/O: input/output
LC: licking chamber
MAD: Median absolute deviation
ML: Mediolateral
NP: nose poke
OFC: orbitofrontal cortex
PPV: positive predictive values
RL: Reversal learning
RA: Reward anticipation
SG: strain gauge
TS: touch sensor
TTL: transistor-transistor logic
TP: True positive
WD: Walking delay

---

**Extended Data Figure 1-1.**
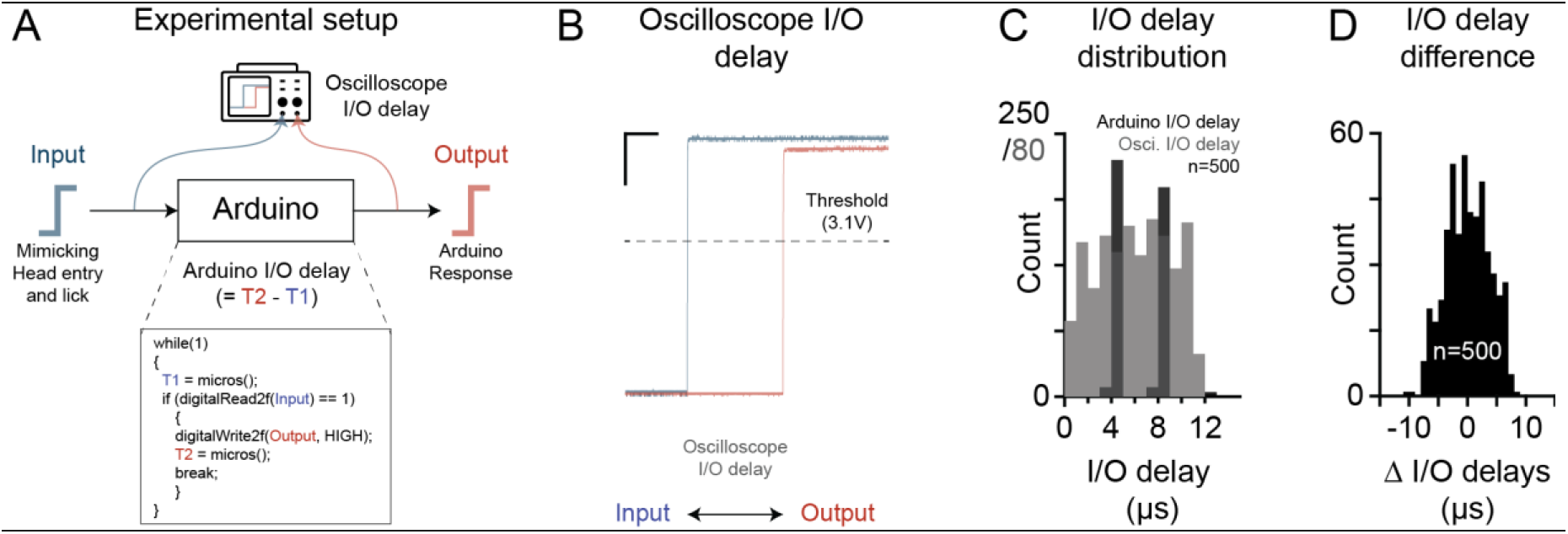
Evaluation of the input/output reactivity of FreiBox. **(A-B)**. Measuring behavior requires integration and control of a myriad of different instruments in response to animal behavior by respecting a sub-millisecond timescale resolution. For example, the inter-lick interval is distributed around 100 ms (Figure 2D), meaning that an Arduino controller has to integrate external information very quickly to measure animal behavior. To quantify the speed of FreiBox to integrate, respond, and timestamp such digital inputs, we conducted an experiment to measure the response reactivity (‘I/O delay’) of an Arduino Mega to respond to an incoming TTL input. **(A)**. Experimental setup and Arduino code used to measure the I/O delay of an Arduino Mega (‘Output’) receiving an incoming TTL input (‘Input’) driven by an experimenter controlling a 5V manual-push button. Arduino Mega was programmed to register the timestamp of the Input TTL (T1), detected with the Arduino library ‘DIO2’, and the timestamp of the TTL output (T2) right after its emission. In parallel, an oscilloscope (DSO-1204e, Voltcraft) connected to a computer (software: DSO3104) recorded both input and output TTL to quantify the I/O delay at a fast-sampling (500 kHz). **(B)**. Example of the TTL traces recorded with an oscilloscope (sampling rate 500 kHz, scales: 5 μs, 1V). The oscilloscope I/O delay was obtained offline by measuring the delay between the Input and Output TTLs when their voltages were crossing 3.1 V (dashed line). **(C-D)**. In this experiment, we used 2 different boxes and mimicked 250 TTL inputs in each box by pressing a push button (n=500 total trials). For these boxes, the I/O delay recorded with the Arduino (Box1 vs Box2, 6.03×10^−6^ ± 1.35×10^−7^ s vs 5.86×10^−6^ ± 1.28×10^−7^ s, n=250, Wilcoxon signed rank test, Z=−0.711, *p*=0.478) and the oscilloscope (Box1 vs Box2, 5.90×10^−6^ ± 2.01×10^−7^ s vs 6.13×10^−6^ ± 1.94×10^−7^ s, n=250, paired t-test, T=−0.862, *p*=0.390) were not statistically different. Hence, we pooled the data together. **(C)**. Histogram distribution showing that the Arduino and Oscilloscope I/O delays are not statistically different (‘Arduino I/O delay’ vs ‘Osci. I/O delay’, 6.01×10^−6^ ± 1.40×10^− 7^ s vs 5.94×10^−6^ ± 9.30×10^−8^ s, n=500, Friedman Repeated Measures Analysis of Variance on Ranks, X^2^= 8×10^−3^, *p*=0.929). This result illustrates that an Arduino Mega 2560 can react to an incoming input with a delay less than 20 μs. **(D)**. Histogram distribution of the delay difference recorded simultaneously with the Arduino and the oscilloscope, showing that an Arduino can also register 2 close events with a precision of ±10 μs.

**Extended Data Figure 1-2.**
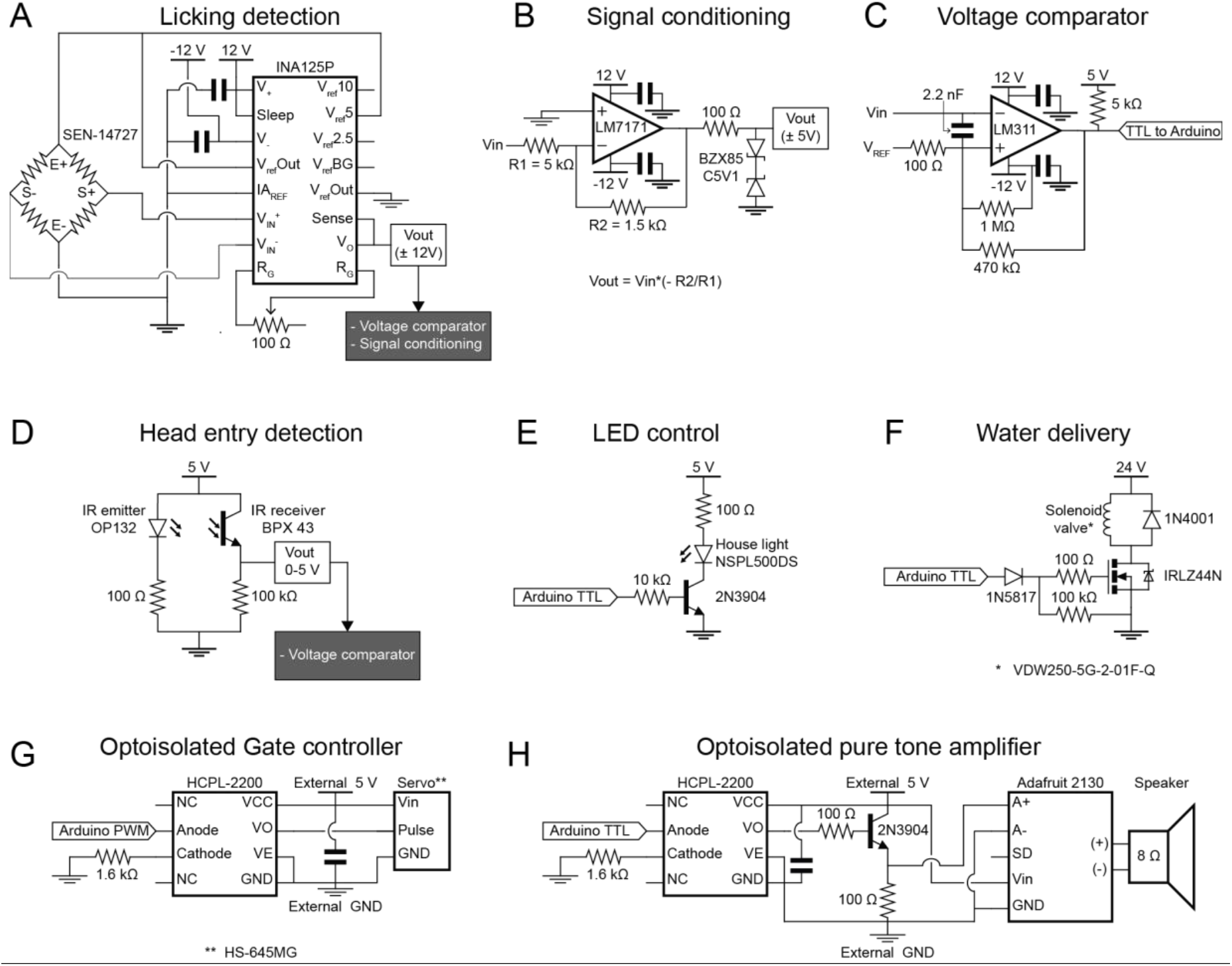
Electronic circuits of the FreiBox modules. **(A-B)**. Amplification (A) and conditioning (B) circuit of the SG signal. The signal conditioning circuit is used to divide the voltage of the amplified signal (±12V) in a range that is acceptable for recording with most of DAC board (±5V). Note that 100Ω trimmer in (A) is used to adjust the sensitivity of the instrumental amplifier. Voltage comparator circuit used to generate a TTL when the input voltage (Vin) exceeds the value of a reference voltage (Vref). This circuit is used to detect both individual licks, and the IR beam breaking induced by a head entry in the NP or LC. The Vref is controlled by a precise potentiometer (10KΩ, tolerance 5%, 10 turns) to fine-tune the licking detection or by a trimmer for head entry detection. Electronic circuit of the head entry detection system. The generated signal (Vout) is sent to the voltage comparator circuit described in (C) to generate TTL signal during a NP. **(E-H)**. Electronic circuits of the Light-house LED controller (E), water delivery system (F), the optoisolated gate controller (G) and the optoisolated auditory cure amplifier (H). Optoisolators are used to reduce the electrical noise induced by the motor in the FreiBox circuit and in the speaker.

**Extended Data Figure 2-1.**
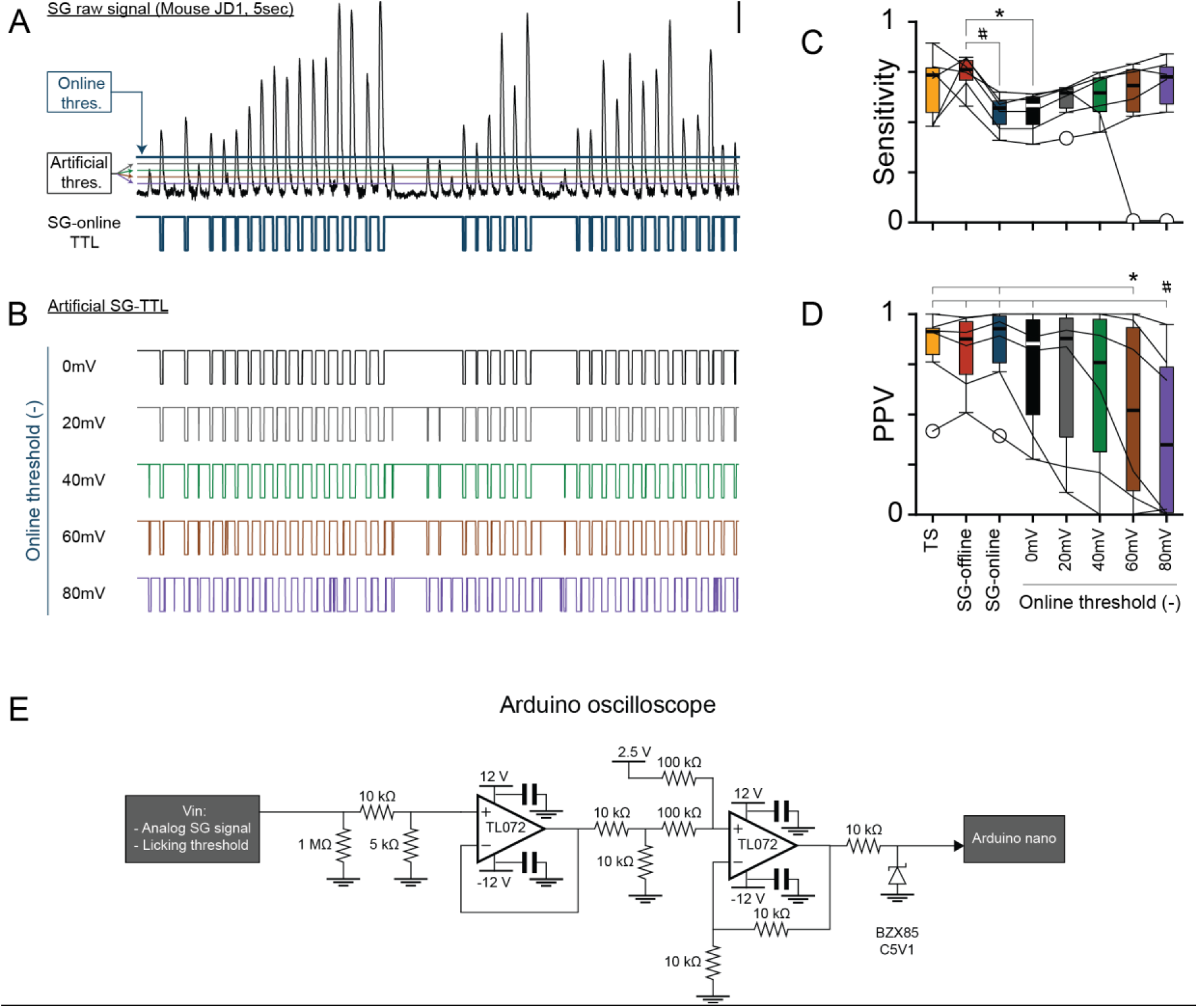
Improving online detection of the SG-lickometer. **(A-B)**. A 5 s example of the SG raw signal (A, scale: 0.1 V) and its online detection signal (SG-online TTL) recorded from another mouse (#JD1) than in Fig. 2C. The SG-online TTL is generated when the SGs crosses an online reference voltage threshold set 100 mV above the baseline (‘Online thres.’). By decreasing artificially this reference voltage (A, ‘Artificial thresh.’), we generated multiple artificial SG-TTL traces (B) to evaluate the effect of the voltage threshold on the sensitivity (C) and the PPV (D). Box-and-whisker plots comparing the sensitivity of SG lickometer and of the artificial SG-TTL traces as described in (B) (Friedman repeated measures ANOVA on ranks, n=6, X^2^= 21.643, 7 df, *p*=0.003; TS vs SG-offline vs SG-online vs 0 mV vs 20 mV vs 40 mV vs 60 mV; 0.685 ± 0.0676 vs 0.740 ± 0.0374 vs 0.546 ± 0.0372 vs 0.549 ± 0.0413 vs 0.600 ± 0.0406 vs 0.626 ± 0.0486 vs 0.577 ± 0.121 vs 0.606 ± 0.126, respectively; describe data: mean ± SEM). * and #, Tukey test post hoc, q=5, *p*<0.05. Box-and-whisker plots comparing the PPV of SG lickometer and of the artificial SG-TTL traces as described in (B) (one-way Anova repeated measures, n=6, F=5.523, *p*<0.001; TS vs SG-offline vs SG-online vs 0 mV vs 20 mV vs 40 mV vs 60 mV; 0.823 ± 0,0873 vs 0.813 ± 0.0785 vs 0.827 ± 0.0972 vs 0.729 ± 0.129 vs 0.684 ± 0.164 vs 0.621 ± 0.175 vs 0.517 ± 0.189 vs 0.402 ± 0.178, respectively; describe data: mean ± SEM # Tukey post hoc test; 80 mv vs TS, 80 mV vs SG-Offline, 80 mV vs SG-Online, 80 mV vs 0 mV; q=86.365, 86.205, 86.417, 84.941, respectively; *p*=0.002, 0.002, 0.002, 0.026 respectively. * Tukey post hoc; 60 mV vs TS and 60 mV vs SG Online; q=84,630 and 84.683 respectively; p=0.044 and 0.04, respectively.

**Extended Data Figure 3-1.**
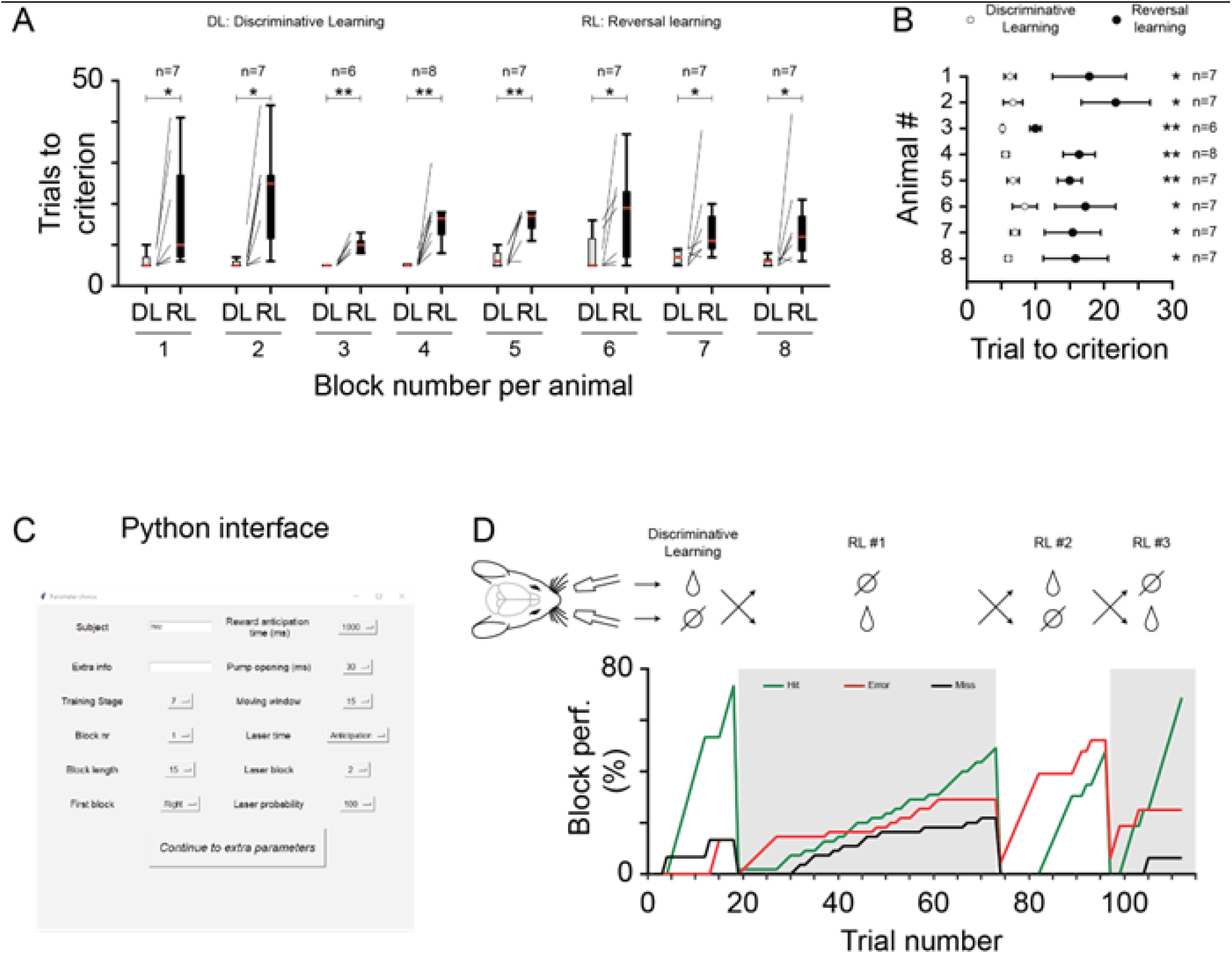
Individual behavioral data and serial reversal learning setting. **(A-B)**. Number (A) and average (B, mean ± SEM) of trials to reach the criterion of each individual mouse used in the figure 4. For the mice 1 (n=7, t=−2.492, *p*=0.047), 2 (n=7, t= −3.739, *p*=0.01), 3 (n=6, t = −6.100, *p*=0.002), 4 (n=8, t= −4.633, *p*=0.002), and 5 (n=8, t= −4.757, *p*=0.003), a paired T-test was performed to compare the numbers of trials to criterion during discriminative learning (DL) and reversal learning (RL) blocks. In contrast, Wilcoxon signed rank tests were performed for the mice 6 (n=7, z= 2.201, *p*=0.031), 7 (n=7, t = 2.201, *p*=0.0031) and 8 (n=7, t = 2.197, *p*=0.0031). The * and ** show significance at the risk α=0.05 and 0.01 respectively. **(C-D)**. By changing the number of blocks in the GUI interface (C), we are able to control a serial intrasession RL in well trained Thy1-GCamP6f mice as shown in D. OFC GCamP6f expression as well as Ca^2+^ imaging recorded during the same behavioral session are shown in Supplement S4.

**Extended Data Figure 4-1.**
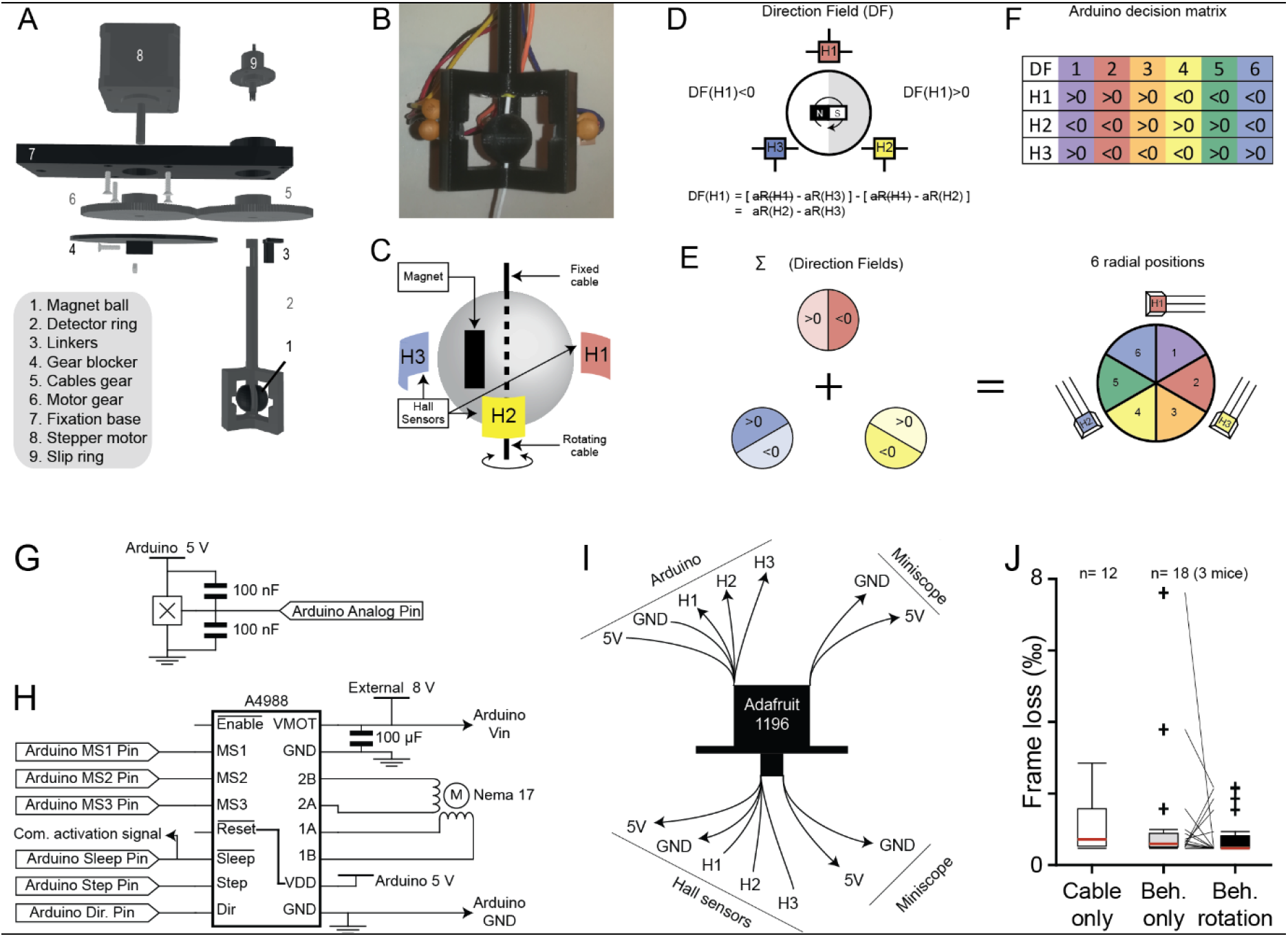
The FreiBox commutator. **(A-C)**. Neurophysiological recordings in freely moving animals require untangling of the miniscope coaxial cables connected to the freely moving implanted animal. To solve that problem, previous designs developed a well-established method to detect the rotation of a cable implant by tracking the relative position of a magnet to a sensor called Hall sensor (Barbera et al., 2020; Fee & Leonardo, 2001; Liberti et al., 2017). By modifying a commutator from the Gardner’s lab (Liberti et al., 2017), we designed a low-cost (111 €) open-source commutator as described in the exploded view (A). By adding a magnetic ring rotary encoder (B), we improved the detection of the cable position and constrained the cable movement. This design offers the advantage of maintaining the magnet close to the detector ring even when the connected mouse moves in a spacious environment. The schematic representation of the ring commutator (C) shows that a magnet glued to the miniscope coaxial cable can rotate along the cable axis between 3 fixed Hall sensors arranged in a circle around the cable (H1, H2 and H3). **(D-F)**. To track the position of the magnet into the ring detector, the direction field (DF) of each Hall sensor is calculated as described in (D) by an Arduino using the function analogRead (‘aR’). By combining the direction fields of the 3 Hall sensors (E), it is possible to define the orientation of the magnet in the field forms by the 3 Hall sensors (among 6 radial positions), and adjust the cable position on the basis of a decision matrix (F). **(G-I)**. To build the FreiBox commutator, the 3 Hall sensors (G) and a motor driver card (H) are connected to the step motor and the Arduino. The signals from the miniscope and the Hall sensors are relayed via a slip ring (I). **(J)**. To test whether the commutator rotation induces additional data loss during miniscope recordings, we compared the percent of frame loss when the miniscope was connected to the DAC board with a simple coaxial cable and left immobile on a table (‘Cable only’, n=12 sessions), to acquisitions performed during behavioral sessions (18 sessions, n=3 mice). More precisely, we calculated the frame loss percent when the commutator was actively rotating (‘Beh. Rotation’) or not (‘Beh. Only’) during imaging sessions in mice performing the task. As illustrated in the box-and-whisker plots (J), we found that the basal percent of frame loss (‘Cable only’) did not statistically increase when the signal is transmitted via an inactive commutator (‘Behavioral Only’ vs ‘Cable only’, 0.806 ± 0.444 vs 0.715 ± 0.238, n=18 vs 12, Mann-Whitney Rank Sum Test, U=88.00, *p*= 0,406), nor during active rotation (‘Behavioral rotation’ vs ‘Cable only’, n=18 vs 12, Mann-Whitney Rank Sum Test, U=66.500, *p*=0,060), nor when active rotation is compared to inactive condition (‘Behavioral Rotation’ vs ‘Behavioral only’, 0.371 ± 0.158 vs 0.715 ± 0.238, n=18, Wilcoxon signed rank test, Z=−0.973, p=0.358). **References.** Barbera, G., Zhang, Y., Werner, C., Liang, B., Li, Y., & Lin, D. T. (2020). An open source motorized swivel for in vivo neural and behavioral recordings. MethodsX, 7, 101167. https://doi.org/10.1016/J.MEX.2020.101167 Fee, M. S., & Leonardo, A. (2001). Miniature motorized microdrive and commutator system for chronic neural recording in small animals. Journal of Neuroscience Methods, 112(2), 83–94. https://doi.org/10.1016/S0165-0270(01)00426-5 Liberti, W. A., Perkins, L. N., Leman, D. P., & Gardner, T. J. (2017). An open source, wireless capable miniature microscope system. Journal of Neural Engineering, 14(4), 045001. https://doi.org/10.1088/1741-2552/AA6806

**Extended Data Figure 4-3.**
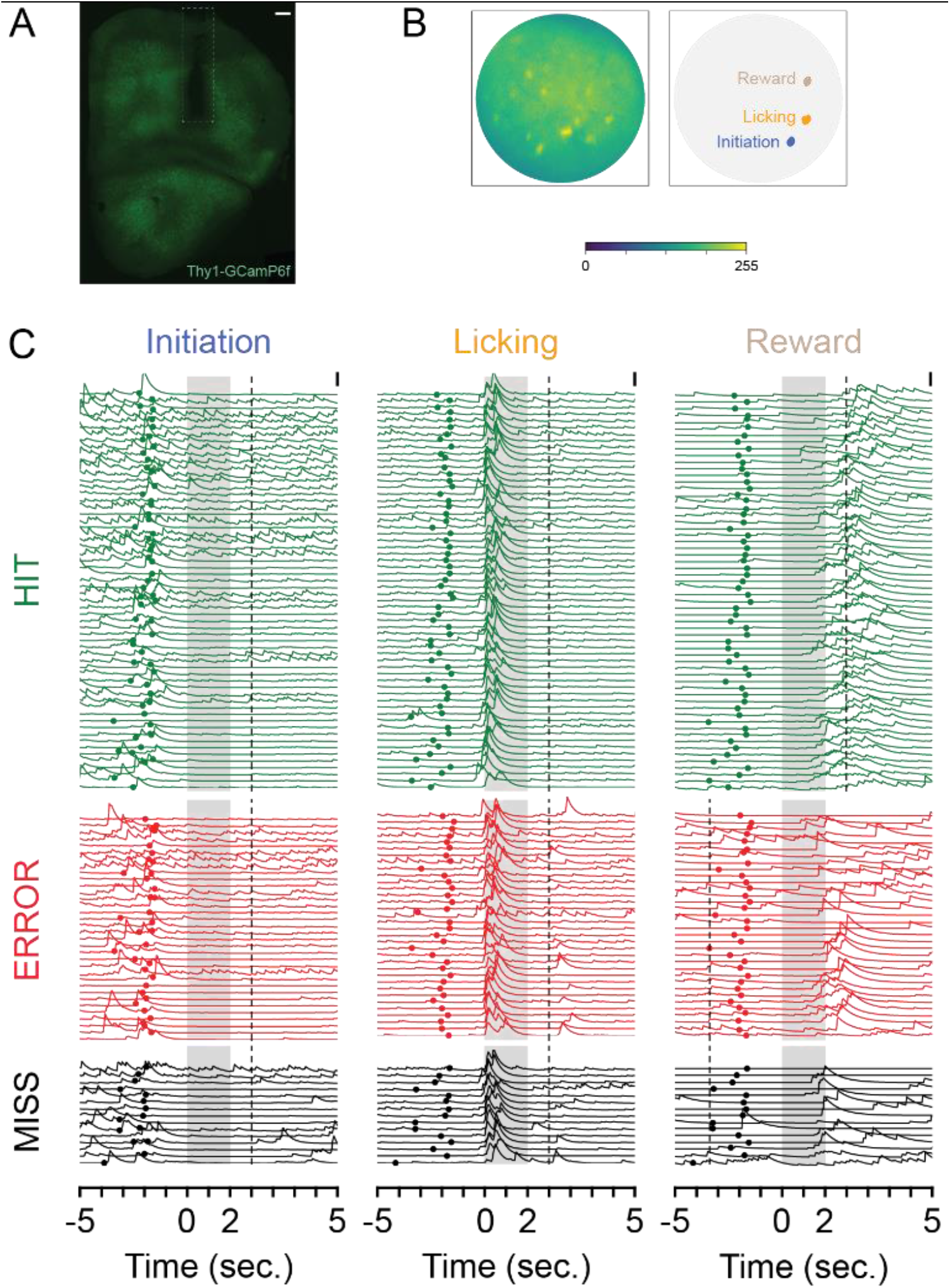
OFC calcium imaging during intra-session RL. **(A)**. Thy1-GCaMP6f expression and GRIN lens track in OFC (scale: 250μm). **(B-C)**. Examples of 3 modulated neurons (B) and their calcium traces (C) aligned to the head entry in the LC (0 s), during the same behavioral session as shown in Supplement S3D. The Maximum projection picture (B left) was obtained on the first 10 k frames of the recording session. The spatial footprints (B right) and the normalized calcium traces (C) were obtained with the open-source pipeline CalmAn (gSig/gSiz: 3/13). The calcium traces represent the normalized denoised temporal traces (CalmAn parameter C, scales: 5 normalized df) and were colored in accordance with their spatial footprints shown in B right. Normalized calcium traces were aligned to the beginning of the LR windows (shaded in grey), sorted by trial type (hit, error, or miss) and their trial number (shown here in an ascending order). The colored dots indicate the nose poke timestamps. For the hit trials, the dashed lines indicate the end of the anticipatory response windows, immediately followed by a water reward delivery. A precise list of the components and providers is available on https://github.com/Optophys-Lab/FreiBox. The prices have been updated on June 2022.

## REFERENCES

Aharoni, D., Khakh, B. S., Silva, A. J., & Golshani, P. (2019). All the light that we can see: a new era in miniaturized microscopy. Nature Methods, 16(1), 11–13. https://doi.org/10.1038/S41592-018-0266-X

Bollu, T., Ito, B. S., Whitehead, S. C., Kardon, B., Redd, J., Liu, M. H., & Goldberg, J. H. (2021). Cortex-dependent corrections as the tongue reaches for and misses targets. Nature, 594(7861), 82–87. https://doi.org/10.1038/S41586-021-03561-9

Boulougouris, V., Dalley, J. W., & Robbins, T. W. (2007). Effects of orbitofrontal, infralimbic and prelimbic cortical lesions on serial spatial reversal learning in the rat. Behavioural Brain Research, 179(2), 219–228. https://doi.org/10.1016/J.BBR.2007.02.005

Brady, A. M., & Floresco, S. B. (2015). Operant procedures for assessing behavioral flexibility in rats. Journal of Visualized Experiments, 96. https://doi.org/10.3791/52387

Castañé Anna, A., Theobald, D. E. H., & Robbins, T. W. (2010). Selective lesions of the dorsomedial striatum impair serial spatial reversal learning in rats. Behavioural Brain Research, 210(1), 74–83. https://doi.org/10.1016/J.BBR.2010.02.017

Catanese, J., & Jaeger, D. (2021). Premotor Ramping of Thalamic Neuronal Activity Is Modulated by Nigral Inputs and Contributes to Control the Timing of Action Release. Journal of Neuroscience, 41(9), 1878–1891. https://doi.org/10.1523/JNEUROSCI.1204-20.2020

Chen, T. W., Li, N., Daie, K., & Svoboda, K. (2017). A Map of Anticipatory Activity in Mouse Motor Cortex. Neuron, 94(4), 866-879.e4. https://doi.org/10.1016/J.NEURON.2017.05.005

Crompe, B. de la, Aristieta, A., Leblois, A., Elsherbiny, S., Boraud, T., & Mallet, N. P. (2020). The globus pallidus orchestrates abnormal network dynamics in a model of Parkinsonism. Nature Communications, 11(1). https://doi.org/10.1038/S41467-020-15352-3

Floresco, S. B., & Jentsch, J. D. (2011). Pharmacological enhancement of memory and executive functioning in laboratory animals. Neuropsychopharmacology: Official Publication of the American College of Neuropsychopharmacology, 36(1), 227–250. https://doi.org/10.1038/NPP.2010.158

Forman, C. J., Tomes, H., Mbobo, B., Burman, R. J., Jacobs, M., Baden, T., & Raimondo, J. v. (2017). Openspritzer: an open hardware pressure ejection system for reliably delivering picolitre volumes. Scientific Reports, 7(1). https://doi.org/10.1038/S41598-017-02301-2

Galiñanes, G. L., Bonardi, C., & Huber, D. (2018). Directional Reaching for Water as a Cortex-Dependent Behavioral Framework for Mice. Cell Reports, 22(10), 2767–2783. https://doi.org/10.1016/J.CELREP.2018.02.042

Giovannucci, A., Friedrich, J., Gunn, P., Kalfon, J., Brown, B. L., Koay, S. A., Taxidis, J., Najafi, F., Gauthier, J. L., Zhou, P., Khakh, B. S., Tank, D. W., Chklovskii, D. B., & Pnevmatikakis, E. A. (2019). CaImAn an open source tool for scalable calcium imaging data analysis. ELife, 8. https://doi.org/10.7554/ELIFE.38173

Goltstein, P. M., Reinert, S., Glas, A., Bonhoeffer, T., & Hübener, M. (2018). Food and water restriction lead to differential learning behaviors in a head-fixed two-choice visual discrimination task for mice. PLOS ONE, 13(9), e0204066. https://doi.org/10.1371/JOURNAL.PONE.0204066

Guo, Z. v., Inagaki, H. K., Daie, K., Druckmann, S., Gerfen, C. R., & Svoboda, K. (2017). Maintenance of persistent activity in a frontal thalamocortical loop. Nature 2017 545:7653, 545(7653), 181–186. https://doi.org/10.1038/nature22324

Histed, M. H., Pasupathy, A., & Miller, E. K. (2009). Learning substrates in the primate prefrontal cortex and striatum: sustained activity related to successful actions. Neuron, 63(2), 244–253. https://doi.org/10.1016/J.NEURON.2009.06.019

Isett, B. R., Feasel, S. H., Lane, M. A., & Feldman, D. E. (2018). Slip-Based Coding of Local Shape and Texture in Mouse S1. Neuron, 97(2), 418–433.e5. https://doi.org/10.1016/j.neuron.2017.12.021

Izquierdo, A., Brigman, J. L., Radke, A. K., Rudebeck, P. H., & Holmes, A. (2017). The neural basis of reversal learning: An updated perspective. Neuroscience, 345(March), 12–26. https://doi.org/10.1016/j.neuroscience.2016.03.021

Mayrhofer, J. M., El-Boustani, S., Foustoukos, G., Auffret, M., Tamura, K., & Petersen, C. C. H. (2019). Distinct Contributions of Whisker Sensory Cortex and Tongue-Jaw Motor Cortex in a Goal-Directed Sensorimotor Transformation. Neuron, 103(6), 1034–1043.e5. https://doi.org/10.1016/J.NEURON.2019.07.008

Morrissette, A. E., Chen, P. H., Bhamani, C., Borden, P. Y., Waiblinger, C., Stanley, G. B., & Jaeger, D. (2019). Unilateral optogenetic inhibition and excitation of basal ganglia output affect directional lick choices and movement initiation in mice. Neuroscience, 423, 55. https://doi.org/10.1016/J.NEUROSCIENCE.2019.10.031

Parker, N. F., Cameron, C. M., Taliaferro, J. P., Lee, J., Choi, J. Y., Davidson, T. J., Daw, N. D., & Witten, I. B. (2016). Reward and choice encoding in terminals of midbrain dopamine neurons depends on striatal target. Nature Neuroscience, 19(6), 845–854. https://doi.org/10.1038/NN.4287

Pasupathy, A., & Miller, E. K. (2005). Different time courses of learning-related activity in the prefrontal cortex and striatum. Nature, 433(7028), 873–876. https://doi.org/10.1038/NATURE03287

Peng, T., Thorn, K., Schroeder, T., Wang, L., Theis, F. J., Marr, C., & Navab, N. (2017). A BaSiC tool for background and shading correction of optical microscopy images. Nature Communications 2017 8:1, 8(1), 1–7. https://doi.org/10.1038/ncomms14836

Schoenbaum, G., Garmon, J. W., & Setlow, B. (2001). A novel method for detecting licking behavior during recording of electrophysiological signals from the brain. Journal of Neuroscience Methods, 106(2), 139–146. https://doi.org/10.1016/S0165-0270(01)00341-7

Schoenbaum, G., Roesch, M. R., Stalnaker, T. A., & Takahashi, Y. K. (2009). A new perspective on the role of the orbitofrontal cortex in adaptive behaviour. Nature Reviews. Neuroscience, 10(12), 885–892. https://doi.org/10.1038/NRN2753

Siegle, J. H., López, A. C., Patel, Y. A., Abramov, K., Ohayon, S., & Voigts, J. (2017). Open Ephys: an open-source, plugin-based platform for multichannel electrophysiology. Journal of Neural Engineering, 14(4). https://doi.org/10.1088/1741-2552/AA5EEA

Wang, G., & Fowler, S. C. (1999). Effects of haloperidol and clozapine on tongue dynamics during licking in CD-1, BALB/c and C57BL/6 mice. Psychopharmacology 1999 147:1, 147(1), 38–45. https://doi.org/10.1007/S002130051140

Wang, Y., Yin, X., Zhang, Z., Li, J., Zhao, W., & Guo, Z. v. (2021). A cortico-basal ganglia-thalamo-cortical channel underlying short-term memory. Neuron, 109(21), 3486–3499.e7. https://doi.org/10.1016/J.NEURON.2021.08.002

White, S. R., Amarante, L. M., Kravitz, A. v., & Laubach, M. (2019). The Future Is Open: Open-Source Tools for Behavioral Neuroscience Research. ENeuro, 6(4). https://doi.org/10.1523/ENEURO.0223-19.2019

Wu, Z., Litwin-Kumar, A., Shamash, P., Taylor, A., Axel, R., & Shadlen, M. N. (2020a). Context-Dependent Decision Making in a Premotor Circuit. Neuron, 106(2), 316–328.e6. https://doi.org/10.1016/J.NEURON.2020.01.034

Wu, Z., Litwin-Kumar, A., Shamash, P., Taylor, A., Axel, R., & Shadlen, M. N. (2020b). Context-Dependent Decision Making in a Premotor Circuit. Neuron, 106(2), 316–328.e6. https://doi.org/10.1016/J.NEURON.2020.01.034

